# Sexually dimorphic ATF4 expression in the fat confers female stress tolerance in *Drosophila melanogaster*

**DOI:** 10.1101/2024.12.27.630478

**Authors:** Lydia Grmai, Melissa Mychalczuk, Aditya Arkalgud, Deepika Vasudevan

**Affiliations:** Department of Cell Biology, University of Pittsburgh School of Medicine, Pittsburgh, PA, USA

**Keywords:** ATF4, transformer, adipose, dimorphism, splicing, doublesex, methioninase, stress

## Abstract

Metabolic differences between males and females have been well documented across many species. However, the molecular basis of these differences and how they impact tolerance to nutrient deprivation is still under investigation. In this work, we use *Drosophila melanogaster* to demonstrate that sex-specific differences in fat tissue metabolism are driven, in part, by dimorphic expression of the Integrated Stress Response (ISR) transcription factor, ATF4. We found that female fat tissues have higher ATF4 activity than their male counter parts under homeostatic conditions. This dimorphism was partly due to a female bias in transcript abundance of specific *ATF4* splice isoforms. We found that the canonical sex determinants *transformer* (*tra*) and *doublesex* (*dsx*) drive such dimorphic *ATF4* transcript abundance. These differences persist in a genetic model of nutrient deprivation, where female animals showed greater resistance to lethality than males in an ATF4-dependent manner. These results suggest that higher ATF4 activity confers higher tolerance to stress in females. Together, our data describe a previously unknown facet of ISR signaling wherein sexual identity of adipose tissue confers differential stress tolerance in males and females. Since energy storage mechanisms are known to be dimorphic and have been linked to ATF4 regulation, our studies provide a mechanistic starting point for understanding how sexual identity influences metabolic disease outcomes.

## Introduction

Sex differences in metabolism have been well documented across many species^1^, and several of these differences are ascribed to the sexually dimorphic nature of adipose tissue^2,3^. Population studies across geographies have demonstrated that females bear more adipose tissue and total higher body fat than men^4–6^. The clinical ramifications of such dimorphic adipose tissue biology include substantial sex differences in susceptibility to metabolic diseases and cardiovascular disorders^7^, though females and males show different vulnerabilities to different metabolic stressors. For example, as of 2018 (national Center for Health Statistics, Center for Disease Control) the female incidence of severe obesity was 67% higher than the male incidence^8,9^, which has been attributed in part to higher efficiency of fat storage mechanisms in females compared with males^10^. However, males show a higher predisposition to diabetes and insulin resistance^11^. Obesity has also been shown to be coincident with, or a comorbidity, for multiple metabolic disorders, including diabetes, polycystic ovarian syndrome, and cardiovascular disease^12–14^, with females being especially vulnerable to these comorbidities^15^. While efforts from several groups have made strides in establishing the molecular differences in male and female adipose tissue^16–19^, these clinical findings underscore the importance of understanding the molecular basis of metabolic sex differences, and specifically how sexually dimorphic gene expression influences adipose tissue physiology and metabolic disease.

In the fruit fly *Drosophila melanogaster,* sexual dimorphism in metabolic tissues impacts development and various aspects of physiology including fecundity, immunity, and lifespan^2,20,21^. As with mammalian sexual dimorphism, metabolic sex differences in *Drosophila* are regulated by the sexual identity of fat tissues. *Drosophila* adipocytes comprise the “fat body”, a highly metabolic organ that performs fat- and liver-like functions. Female identity of *Drosophila* larval adipocytes instructs larger body size due to increased secretion of insulin-like peptides from the brain relative to males^22^. Additionally, sexual identity of the Akh-producing cells in the adult brain (analogous to pancreatic α-cells in mammals^23^) regulates fat metabolism^24,25^. Sex differences in lipid metabolism are also partially dependent on sex differences in abundance of the insect hormone 20-hydroxyecdysone (20E), which primes adult females for higher triglyceride and glycogen stores compared with males^26^.

In *Drosophila*, sex differences in metabolic tissues are directed by effectors of the canonical sex determination pathway^22,27–29^, wherein X chromosome number determines expression of the RNA-binding protein Sex-lethal (Sxl)^30,31^: females (bearing two X chromosomes) express Sxl, whereas males (with one X chromosome) do not. In somatic cells, female-specific Sxl action yields alternative splicing of *transformer (tra)* RNA such that full-length Tra protein is only produced in females^32,33^. Tra binding to mRNA leads to production of female splice variants of the transcripts encoding two key sexual differentiation factors, *doublesex* and *fruitless*^34–37^. Dsx and Fru gene products effect numerous aspects of somatic sexual differentiation, including body size, gonad and genitalia development, gametogenesis, feeding, and courtship behavior^38–42^.

In *Drosophila*, female sexual identity confers higher resistance to a variety of stressors compared with males^43–45^. Canonical sex determinants like Tra and Dsx alter the transcriptional landscape of many somatic tissues^46^, though their effects on mediators of stress response signaling are understudied. In this study, we focus on the Integrated Stress Response (ISR), an evolutionarily conserved signaling pathway that elicits adaptive responses to diverse cellular stressors via the transcription factor ATF4^47^. Loss of ATF4 results in increased susceptibility to stressors such as nutrient deprivation and proteotoxicity^48,49^, among others. Studies in many species, including *Drosophila melanogaster*, have demonstrated that caloric restriction improves lifespan in an ATF4-dependent manner^50–52^; in *Drosophila*, female adult lifespan is more responsive to dietary restriction than male lifespan^53^. Further, highly metabolic tissues such as the fat (‘fat body’ in *Drosophila*) rely upon constitutive activation of ATF4 under physiological conditions^49,51,54,55^ and the sexual identity of the fat body has been reported to modify dimorphic stress tolerance^29^. However, whether ISR/ATF4 activity is dimorphic has not yet been examined.

Here, we show that female *Drosophila* fat body exhibits a higher basal level of ISR activity than the male fat body due to sex-biased expression of *ATF4*. This sex bias is regulated downstream of the canonical sex determinants Tra and Dsx. Further, we find that the female-bias in resistance to nutrient deprivation is at least partly mediated by dimorphic ATF4 expression in the fat body. Summarily, our findings suggest that increased activation of ISR signaling in female fat tissues confers a health and/or survival advantage. By uncovering this new facet of ISR signaling, our work establishes a framework for understanding the molecular links between sexual identity and stress tolerance.

## Results

### Homeostatic ATF4 signaling is sexually dimorphic in larval fat tissues

We and others have previously reported homeostatic ATF4 activity in the third-instar larval fat body^48,51,56^. To determine whether this is sex-biased, we utilized an enhancer trap line, *Thor-lacZ*, which we have previously shown to be reflective of ATF4 transcriptional activity^51,57^. We observed 85% higher β-galactosidase abundance in female wandering third-instar fat bodies compared with males at the same stage (**Fig. 1A-C; Fig. S1A**). Since *Thor* (*Drosophila* ortholog of the *4E-BP* gene) expression can also be regulated downstream of the transcription factor FOXO^58,59^, we verified potential dimorphic ATF4 activity in the fat body using a transgenic *in vivo* reporter for ATF4 activity that contains an intronic element from the *Thor* locus (containing two ATF4 binding sites) driving DsRed expression^51^ (*4EBP^intron^-DsRed*). This reporter is demonstrably ATF4-responsive and unaffected by decreased FOXO expression^51^. Consistent with a dimorphism in *Thor-lacZ* expression, we found *4EBP^intron^-DsRed* expression to be 67% higher (p<0.0001) in females compared with males (**Fig. 1D-F; Fig. S1B**). Since DsRed protein produced from this construct was frequently observed in the cytoplasm, we generated another reporter, *4EBP^intron^-GFP*, using the same *4EBP^intron^* element that faithfully drives GFP expression only in the nucleus (**Fig. S1C-D**). Reassuringly, using this reporter we also observed 63% higher *4EBP^intron^* activity (p<0.0001) in female adipocytes compared with male adipocytes (**Fig. S1C-E**).

**Figure 1.**
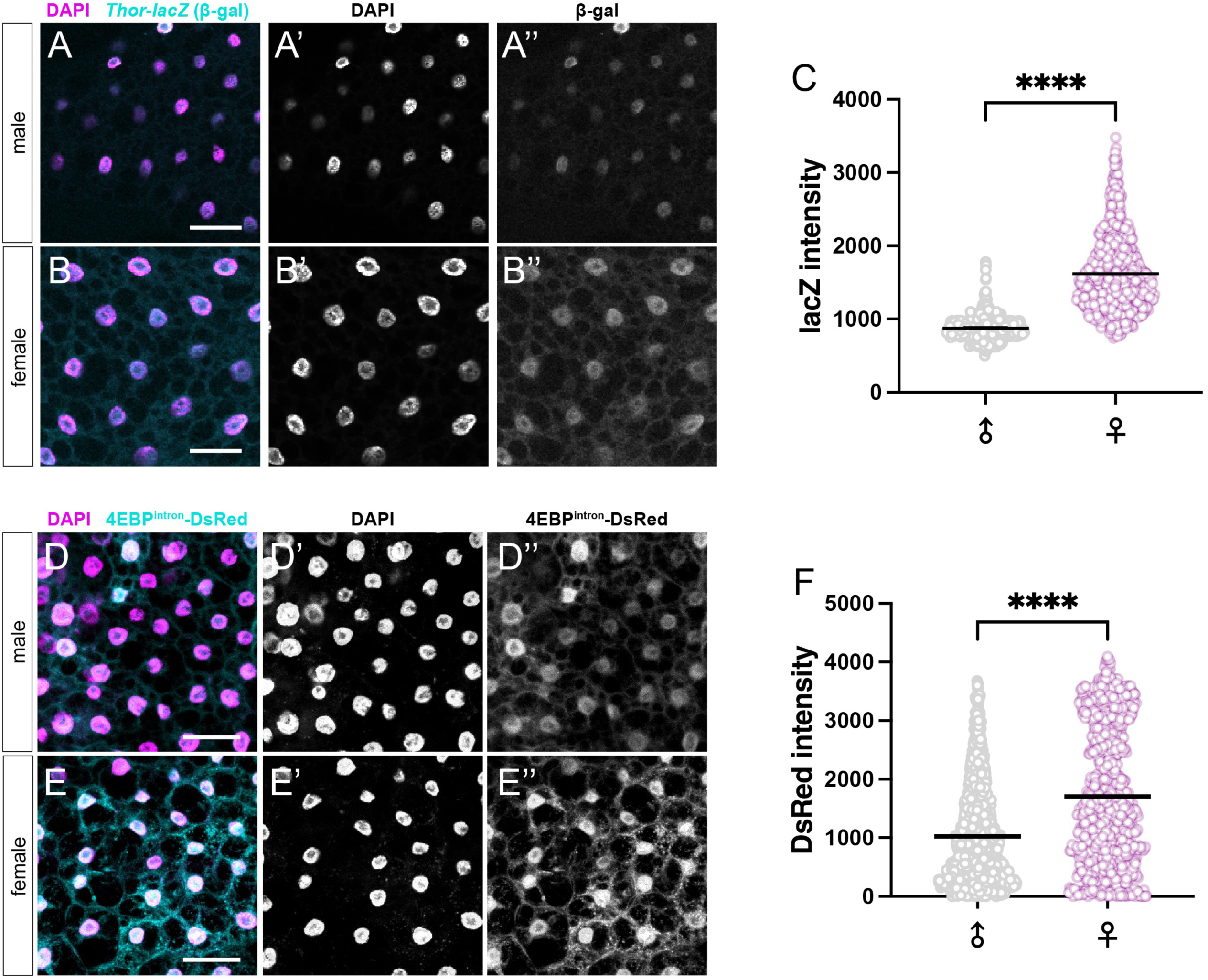
ATF4 activity is sexually dimorphic in larval adipocytes. (A-B) Representative immunofluorescence images of male (A) and female (B) larval fat tissues carrying a *Thor-lacZ* enhancer trap reporter. The inserted *lacZ* gene encodes β-galactosidase (β-gal) gene and carries a nuclear localization signal. In these animals, β-gal abundance is a proxy for *Thor* expression, which appears higher in female adipocytes (B’’) than male adipocytes (A’’). (C) Quantification of nuclear β-gal fluorescence intensity in A-B. (D-E) Representative immunofluorescence images of male (D) and female (E) larval fat tissues carrying a transgenic *4EBP^intron^-DsRed* reporter. (F) Quantification of nuclear DsRed intensity in D-E. Here and throughout the study, experiments were performed on fat body from wandering 3^rd^ instar larvae unless otherwise specified. Here and throughout the study, in all quantifications of confocal images such as C and F, statistical significance was determined using a two-tailed Student’s t-test with Welch’s correction for unequal standard deviations; statistical significance is denoted as follows: *p<0.05; **p<0.01; ***p<0.001; ****p<0.0001. In confocal images, DAPI (magenta) labels nuclei. Scale bars = 50 μm.

### ATF4 (crc) mRNA expression is sexually dimorphic in larval adipocytes in an isoform-specific manner

We reasoned that the higher levels in ATF4 reporter expression observed in female fat body in **Fig. 1 and S1** could be due to differences in *ATF4* mRNA abundance. *Drosophila* ATF4 is encoded by the *cryptocephal* (*crc*) gene; qPCR analyses of fat bodies revealed higher levels of both *Thor* and *crc* mRNA in female fat bodies in comparison to male fat bodies (**Fig. 2A**). There are four annotated *crc* splice isoforms (according to *D.mel* genome annotation, dm6 assembly; **Fig. 2B**): *crc-RA, crc-RB, crc-RE*, and *crc-RF*. The experiment in **Fig. 2A** used primers that detect a common exon-exon junction across all isoforms (**Fig. 2B**, black arrowheads). To examine whether specific *crc* isoforms contributed to dimorphic ATF4 expression, we designed primers across unique exon-exon junctions that allowed us to distinguish the longer isoforms *crc-RA/RE/RF* (**Fig. 2B**, green arrowheads), separately *crc-RE* (which bears a unique exon, **Fig. 2B**, blue arrowheads), and the shortest isoform *crc-RB* (**Fig. 2B**, purple arrowheads). Interestingly, we found a female bias in *crc* isoform abundance using primers that detect *RA/RE/RF* isoforms (**Fig. 2C**), while *crc-RE* primers did not reveal dimorphic abundance (**Fig. 2C**). We also observed dimorphism in *crc-RB* transcript abundance (**Fig. 2C**), indicating that both long- and short-isoforms of ATF4 contribute to dimorphic ISR activity. It is worth mentioning that the only annotated difference between *crc-RA* and *crc*-*RF* is a 23-bp region absent from exon 4 of the *RF* isoform but included in the *RA* isoform (**Fig. 2B**); we were unable to resolve this difference using qPCR or RT-PCR techniques. From these data, we conclude that ATF4 signaling is female-biased in the larval fat body due to greater abundance of *crc-RA/F/B* isoforms in females compared with males.

**Figure 2.**
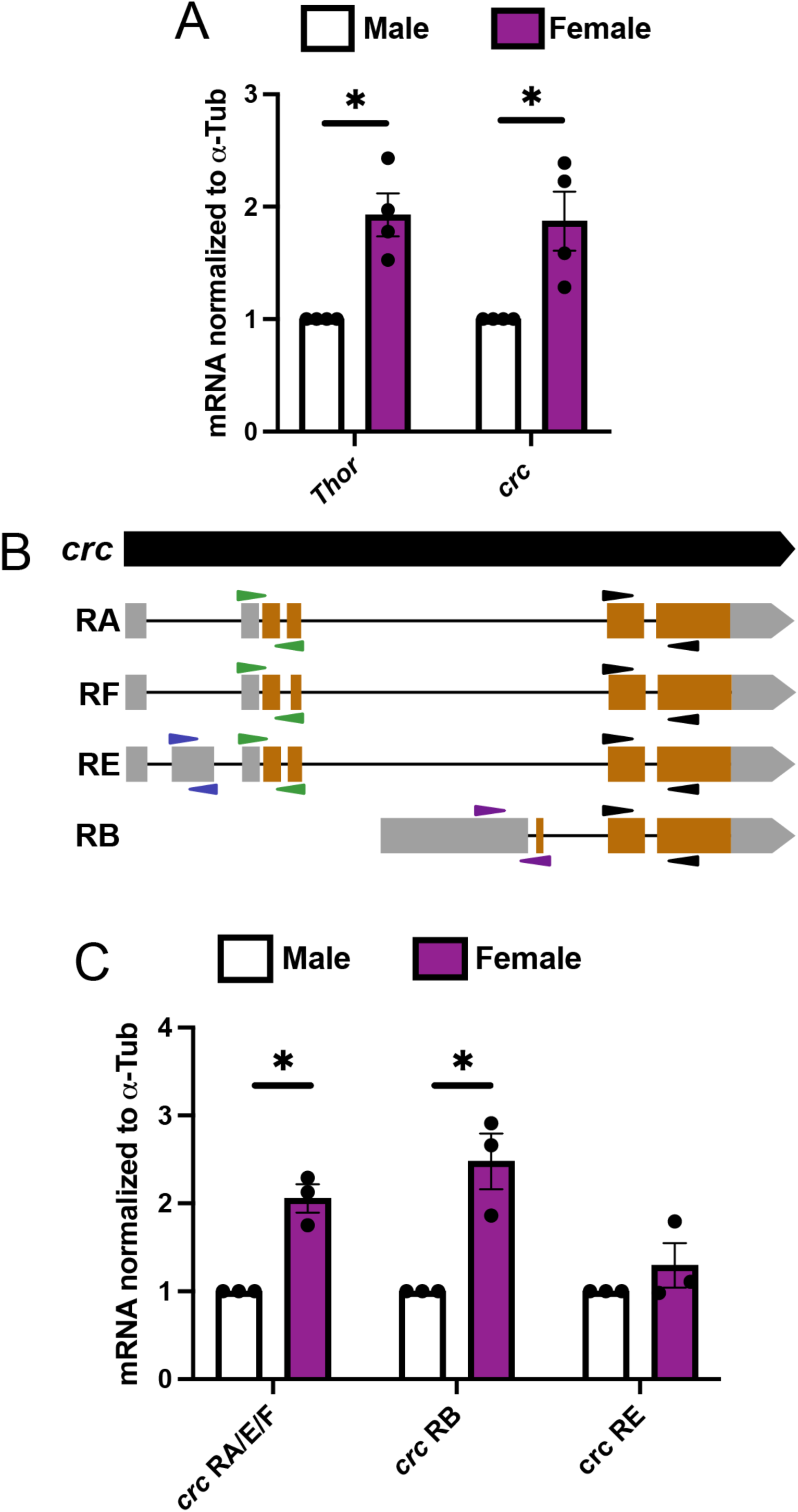
*ATF4* mRNA abundance in larval adipocytes is sexually dimorphic in an isoform-specific manner. (A) mRNA abundance of *Thor* and *crc* in fat bodies from male/female wandering 3^rd^ instar larvae as determined by qRT-PCR. (B) Schematic of *ATF4 (crc)* mRNA isoforms. Orange regions represent ATF4-coding sequences, and gray regions represent all other mRNA exons. Black lines represent intronic regions (exon/intron lengths are not drawn to scale). Pan-isoform primers are indicated with black arrowheads; isoform-specific primers are indicated with blue (*RE*), green (*RA/E/F*), or purple (*RB*) arrowheads. (C) mRNA abundance of specific *crc* isoforms as determined by qRT-PCR. Here and throughout this study, *α-Tub84B* was used as a reference gene for qPCR analyses and statistical significance was determined using ratio-paired two-tailed Student’s t-test.

### Female-biased ATF4 expression is regulated by canonical sex determinants

We next sought to determine whether dimorphic ATF4 expression and activity are established by somatic sex determinants in adipocytes. Canonical sex determination is regulated primarily by the evolutionarily conserved transcription factor Doublesex (Dsx), the founding member of the Doublesex/Mab-3-related transcription factor (DMRT) family^60^. The *dsx* locus encodes two protein isoforms: a male-specific effector (Dsx^M^) and a female-specific effector (Dsx^F^). The cellular decision to express either Dsx isoform is instructed by a splicing cascade downstream of the female-specific RNA-binding proteins Sex-lethal (Sxl)^61^ and Transformer (Tra)^62^. Thus, manipulating *tra* expression elicits sex transformation in a cell-autonomous manner, such that loss of *tra* masculinizes XX cells and over-expression of *tra* results in feminization of XY cells^22,27,40^. We found that homozygous *tra* mutant females (*w^1118^; 4EBP^intron^-GFP/+; tra^1^/tra^KO^*) showed a substantial 59% reduction (p<0.0001) in ATF4 reporter activity in adipocytes compared with adipocytes from control females (*w^1118^; 4EBP^intron^-GFP/+; +*) (**Fig. 3A**). We also examined this effect by depleting *tra* in the female larval fat body (*Dcg-GAL4, UAS-GFP/UAS-tra^RNAi^; 4EBP^intron^-DsRed/+*), which led to a modest 8% reduction (p<0.0001) in *4EBP^intron^-DsRed* expression (**Fig. S2**). We next tested whether *tra* depletion affected *crc* mRNA expression in the fat body. In agreement with our earlier finding that sexually dimorphic ATF4 activity results from higher ATF4 expression in female fat body, we found these tissues also had significantly reduced *crc* transcript abundance upon *tra* depletion (**Fig. 3B**). This reduction was observed for all isoforms we tested (**Fig. 3C-E**), including *crc-RE*, despite the lack of sexual dimorphism in *RE* abundance in control animals. We attribute the differences in the magnitude of effects on *4EBP^intron^-DsRed* versus *crc* mRNA in *Dcg>tra^RNAi^* animals (**Fig. 3A vs B**) to the perdurance of DsRed protein. To test whether the converse is also true (that feminization of male fat body leads to elevated ATF4 activity), we expressed *tra* in male fat body using *Dcg-GAL4*. Indeed, feminization of male fat body (*Dcg-GAL4, UAS-GFP/UAS-tra^F^; 4EBP^intron^-DsRed/+*) more than doubled ATF4 reporter activity (137% increase relative to control male fat body, p<0.0001) (**Fig. 3F, S3A-C**) and nearly doubled *crc* transcript abundance (86% increase relative to control male fat body, p<0.0001) for all isoforms tested (**Fig. 3G-J**).

**Figure 3.**
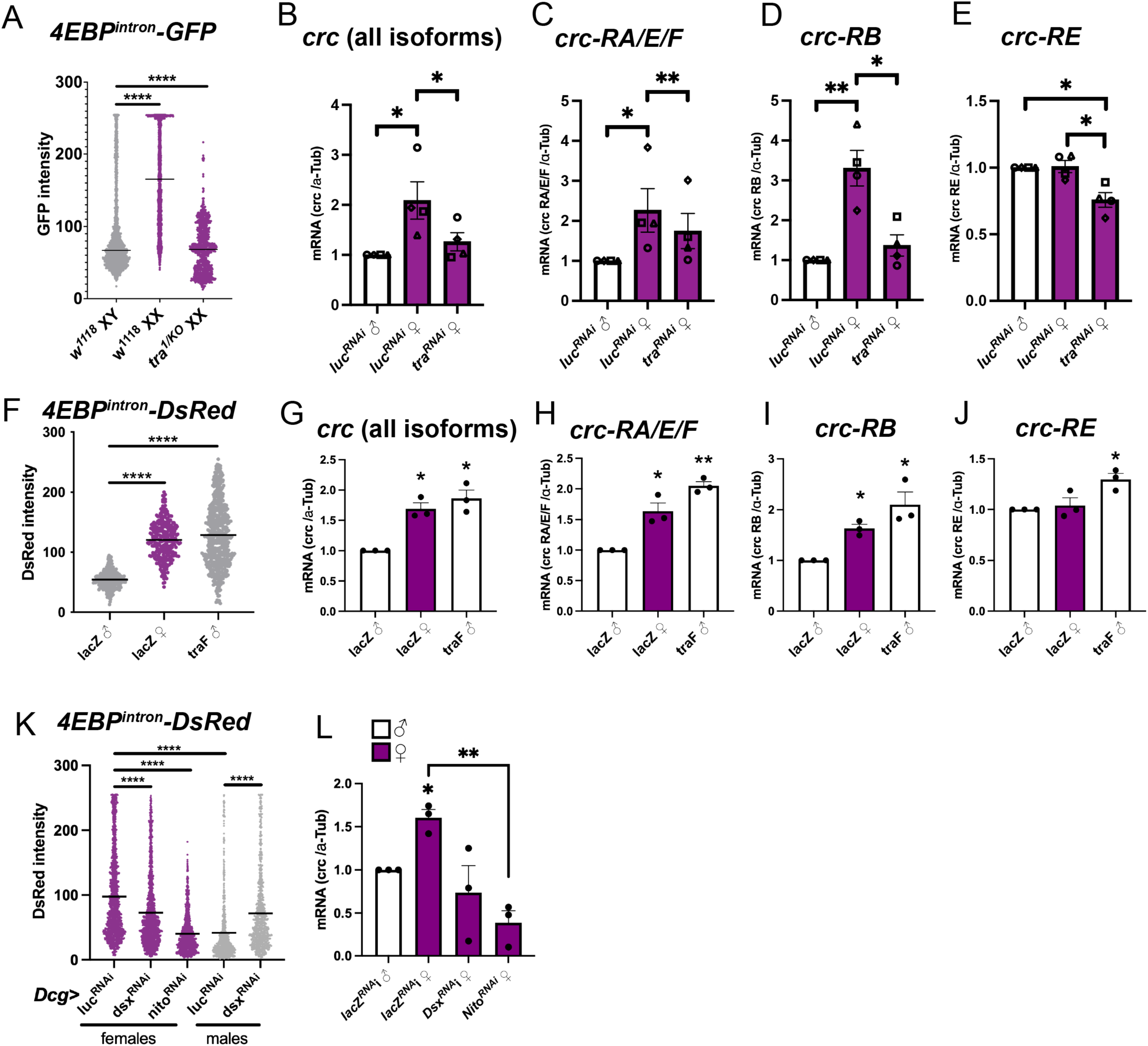
Female sexual identity of larval adipocytes drives higher ATF4 expression in females. (A) Quantification of nuclear *4EBP^intron^-GFP* fluorescence intensity in control (*w^1118^*) or tra-null (*tra^1^*/*tra^KO^*) larval adipocytes. (B-E) qPCR analysis on total and isoform-specific *crc* mRNA abundance in control (*luc^RNAi^*) and genetically masculinized (*tra^RNAi^*) adipocytes. (F) Quantification of nuclear fluorescence intensity of *4EBP^intron^-DsRed* reporter expression in control (*luc^RNAi^*) and genetically feminized (*tra^F^*) larval fat body. (G-J) qPCR analysis on total and isoform-specific *crc* mRNA abundance in tissues from F. (K) Quantification of nuclear fluorescence intensity of *4EBP^intron^-DsRed* reporter expression in control (*luc^RNAi^*) larval fat body and following *dsx* or *nito* knockdown in adipocytes. (L) qPCR analysis on total *crc* mRNA abundance in tissues from K. All experiments in (A-L) utilize the fat body driver *Dcg-GAL4*.

While Dsx is the most well-studied Tra target, there are other effectors of secondary sexual differentiation^22,27^. Thus, we specifically tested whether Dsx also instructs fat body ATF4 expression similar to Tra. The *dsx* gene encodes both male and female Dsx splice isoforms, and loss of *dsx* in either sex is known to produce intersex phenotypes^46,63^. We thus hypothesized that if dimorphic ATF4 expression is regulated by Dsx, then *dsx* knockdown would decrease ATF4 activity in female fat body and increase ATF4 activity in male fat body. Indeed, *dsx* depletion (*Dcg-GAL4, UAS-GFP/UAS-dsx^RNAi^; 4EBP^intron^-DsRed/+*) led to a 25% reduction (p<0.0001) in ATF4 reporter activity in female fat body and a 73% increase (p<0.0001) in ATF4 reporter activity in male fat body (**Fig. 3K, S3D-E, G-H**). This was accompanied by a 54% reduction (p<0.0001) in *crc* transcript abundance following *dsx* knockdown in female fat body (**Fig. 3L**). A comprehensive analysis of Dsx genomic occupancy previously led to the identification of numerous putative direct Dsx transcriptional targets^46^, one of which was *crc*. DamID-seq performed on HA-tagged Dsx^F^ in female fat body revealed occupancy in regulatory regions within the *crc* locus^46^. In addition, bioinformatic analysis using an experimentally determined position weight matrix for Dsx identified multiple high-scoring Dsx binding sequences in the *crc* locus^46^. These analyses present a strong case for direct transcriptional regulation of *crc* by Dsx in the larval fat body.

To further interrogate the sex determinants that govern dimorphic ATF4 activity, we assayed for ATF4 expression and activity upon loss of the RNA-binding protein Spenito (Nito), which effects female sexual identity in the larval fat body^19^ by promoting female splicing of Sxl and Tra^64^ (and via Tra regulation can influence Dsx isoform expression) and. Thus, we sought to test whether dimorphic *crc* expression in fat body is Nito-dependent. We saw that *nito* depletion in female adipocytes (*Dcg-GAL4, UAS-GFP/UAS-nito^RNAi^; 4EBP^intron^-DsRed/+*) caused a 59% reduction (p<0.0001) in ATF4 reporter activity (**Fig. 3K, Fig. S3F**) and a 76% reduction (p<0.0001) in *crc* transcript abundance, similar to the levels observed in control male fat body (**Fig. 3L**). Taken together, our results indicate that sexually dimorphic ATF4 activity in larval fat body is instructed by sexual identity in a cell-autonomous manner, likely via direct transcriptional regulation of *ATF4* by Dsx. However, the difference in the magnitude of DsRed reduction between *tra* mutant versus *Dcg>tra^RNAi^* adipocytes (**Fig. 3A vs S2**) suggest we cannot exclude possible non-autonomous roles for sexual identity in regulating dimorphic ATF4 activity in the fat body.

### Chronic nutrient deprivation stress in fat tissue causes developmental lethality in males

Studies on starvation resistance in *Drosophila* and mammals have revealed that female adipocytes are more tolerant to periods of nutrient scarcity^65,66^. Dietary methionine deprivation has been shown to activate ISR and induce ATF4 expression^47^. Since dietary restriction has systemic effects, we instead used a genetic model of nutrient deprivation in the fat body that allows for cell-autonomous methionine depletion in *Drosophila melanogaster*^67^. In this model, ectopic expression of the bacterial enzyme Methioninase leads to increased catabolism and consequent depletion of the amino acid methionine. Consistent with previous reports of ATF4 reporter induction upon methionine deprivation^51^, we observed that *methioninase* over-expression (*Dcg-GAL4/+; UAS-methioninase/+*) led to higher *Thor* mRNA induction in both male and female adipocytes in comparison to control animals (*Dcg-GAL4/+; UAS-lacZ/+*) (**Fig. S4A**).

Since females are more resilient during periods of nutrient scarcity in *Drosophila*^21,68–70^, we reasoned that inducing *methioninase* expression in the fat during development may differentially impact male and female survival to adulthood. To test this, we performed 24-hour egg lays with equal numbers of age-matched parents and counted the number of adult animals that emerged from each egg lay. Interestingly, *methioninase* expression in the fat body (*Dcg-GAL4, UAS-GFP/+; UAS-methioninase/4EBP^intron^-DsRed*) led to a significantly fewer eclosed males in comparison to control (*Dcg-GAL4, UAS-GFP/+; UAS-lacZ/4EBP^intron^-DsRed*) (**Fig. 4A**). The developmental lethality we observed occurred largely in the pharate stage, immediately preceding eclosion of adults. In contrast, ectopic *methioninase* expression had a much milder effect on the number of eclosed females (**Fig. 4A**), supporting the notion that females are more resilient to nutrient deprivation. As with many GAL4 drivers employed for tissue-specific gene expression^71^, *Dcg-GAL4* expression is not restricted to fat tissues during development or adulthood^72,73^. Thus, we assessed developmental lethality from *methioninase* expression using two other widely employed fat body drivers: *R4-GAL4* and *3.1 Lsp2-GAL4*. We found that *methioninase* expression using either of these fat body drivers phenocopied, to varying degrees, the male developmental lethality observed using *Dcg-GAL4* (**Fig. 4B-C**). Thus, we conclude our developmental lethality phenotypes are primarily due to fat body-autonomous *methioninase* expression.

**Figure 4.**
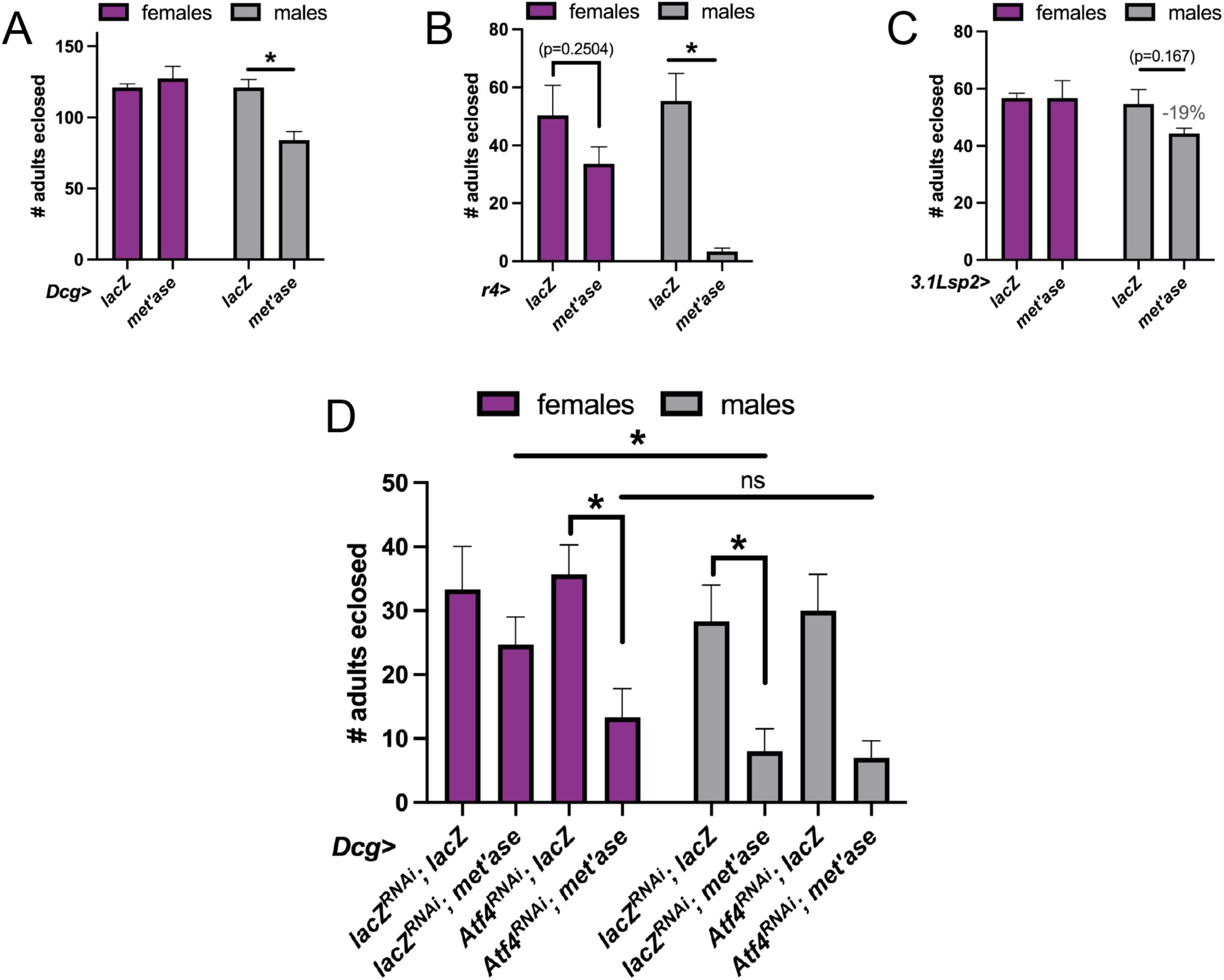
Chronic nutrient deprivation stress causes developmental lethality in males. (A-C) Quantification of males and females eclosed following a 24-hr egg lay from *Dcg-GAL4* (A), *R4-GAL4* (B), or *3.1 Lsp2-GAL4* (C) females crossed to either *UAS-lacZ (lacZ)* or *UAS-methioninase (met’ase)* males. (D) Quantification of males and females eclosed following a 24-hr egg lay from *Dcg-GAL4* females crossed to males of the indicated genotypes (x-axis) to test protective role of ATF4 in preventing female lethality during *methioninase* expression. Statistical significance in A-E was evaluated using a series of unpaired two-tailed Student’s t-tests with Welch’s correction.

In performing genetic nutrient deprivation experiments using *Dcg-GAL4*, we observed that severity of Methioninase-induced developmental lethality increased in subsequent iterations, in that successive 24-hour egg lays performed using the same parents over a three-day period exhibited progressively more severe developmental lethality (**Fig. S4B-D**). We postulate that this is not due to a decline in the fertility of the female parent, since we did not observe a decrease in the total number of progeny produced in the control (*>lacZ*) crosses (**Fig. S4B-D**). These observations led us to question whether “leaky”, GAL4-independent *UAS* expression caused sufficient *methioninase* expression (which all tissues could theoretically experience), to contribute to the observed developmental lethality. To test this, we examined *4EBP^intron^-GFP* activity in animals carrying *UAS-lacZ* or *UAS-methioninase* absent of a GAL4 driver. We were surprised to find that *UAS-methioninase* alone nearly doubled ATF4 reporter activity in both male and female larval fat body compared with *UAS-lacZ* animals (**Fig. S4E**). To assess whether leaky *methioninase* expression contributed to the observed developmental lethality in *Dcg>methioninase* animals, we collected *UAS-lacZ* and *UAS-methioninase* embryos (absent of any *GAL4* transgene) over a 24-hr period and quantified the number of animals that progressed into adulthood. Reassuringly, despite the increased ATF4 activity observed in the presence of *UAS-methioninase*, this increase was not sufficient to cause developmental lethality in males (**Fig. S4F**). Thus, we conclude that GAL4-mediated *methioninase* expression in the fat effected the majority of observed developmental lethality in males in **Fig. 4A-C**.

We next tested whether female sex identity of the fat body and/or higher ATF4 activity therein offer a protective role for females during chronic nutrient deprivation. Since females are more starvation-resistant than males^21,68,70^, we predicted that masculinizing female fat body (via *tra* knockdown) would increase developmental lethality with *methioninase* expression. As shown in **Fig. 4A**, female adult eclosion rate was not significantly affected by *methioninase* expression (**Fig. S4G**, second bar, *Dcg-GAL4/UAS-lacZ^RNAi^;UAS-methioninase/+*) in comparison to control females (**Fig. S4G**, first bar, *Dcg-GAL4/UAS-lacZ^RNAi^;UAS-lacZ/+*). We observed that masculinization of the fat body via *tra* knockdown was sufficient to decrease viability of female animals to adult upon *methioninase* expression (*Dcg-GAL4/UAS-tra^RNAi^;UAS-methioninase/+,* **Fig. S4G**). Finally, we tested whether female resilience to *methioninase* expression relies on higher ATF4 expression. Interestingly, simultaneous depletion of ATF4 and expression of *methioninase* (**Fig. 4D**, *Dcg-GAL4/UAS-ATF4^RNAi^;UAS-methioninase/+*) resulted in a significant increase in female developmental lethality compared with females expressing *methioninase* and a control transgene (*Dcg-GAL4/UAS-ATF4^RNAi^;UAS-lacZ/+*). Such loss of *ATF4* did not appear to further impact the already reduced eclosion rates seen in males with *methioninase* expression (**Fig. 4D**, compare sixth bar to eighth). Taken together, our data support a model wherein high ATF4 activity in female fat body, instructed by cellular sexual identity, confers a protective role to female animals under nutrient deprivation stress.

## Discussion

In this study, we demonstrate that the Integrated Stress Response is sexually dimorphic in larval adipocytes. This dimorphism has implications for sexually dimorphic adipose tissue physiology, wherein the sexual identity of the fat confers a survival advantage to females under nutrient deprivation stress in an ATF4-dependent manner. A growing body of work has established that chromosomally female (XX) animals exhibit higher survival during periods of nutrient scarcity compared with chromosomally male (XY) animals^65,66^. Despite examples of this in both clinical and model organism studies, research is still ongoing to identify the molecular and/or physiological differences between male and female adipocytes that drive sexually dimorphic survival under stress. Our work links previously unreported dimorphic expression of the transcription factor ATF4 in the fat body to survival under nutrient deprivation stress, providing an important molecular clue into the physiology underlying sexually dimorphic stress tolerance.

We used two types of readouts – enhancer-based ATF4 reporter expression and *ATF4* (*crc)* mRNA abundance – to demonstrate that ATF4 expression is female-biased in larval adipocytes (**Fig. 1, S1, 2**). Using tissue-specific loss-of-function experiments, we found that this dimorphism relies on cell-autonomous sexual identity instruction by the canonical sex determinants Tra, Dsx, and Nito (**Fig. 3, S2-3**). Finally, we determined that dimorphic ATF4 expression in the fat underlies sex-biased survival to nutrient deprivation stress: we saw that genetic methionine depletion via *methioninase* expression in the fat body caused developmental lethality disproportionately in males (**Fig. 4A-C, S4B**) and that the survival advantage conferred to females relied both on the sexual identity of adipocytes and on higher basal ATF4 expression therein (**Fig. 4D, S4F**). In summary, our findings implicate fat body ATF4 function in directing sex-specific physiology in *Drosophila melanogaster*. Because the *Drosophila* fat body is analogous to both adipocytes and hepatocytes in mammals, the regulation of ATF4 activity by sexual identity in adipocytes can likely be extended to further understand sexually dimorphic functions of ISR in mammalian fat and liver.

### Sexual dimorphism in metabolic tissues

Increased energy storage capabilities have been implicated in female starvation resistance^68,69,74^, and such sex differences in energy metabolism have been shown and/or presumed to affect long-term disease susceptibilities, such as a higher incidence of obesity in females^8,9,10^ and higher male predisposition to diabetes and insulin resistance^11^. Males and females are differentially sensitive to changes in the abundance of specific energy sources^75–77^ (e.g., carbohydrates versus protein). A clearer understanding of the transcriptional outputs that fuel such sex differences would enable us to better understand energy metabolism in the context of whole organism physiology. Such an understanding might also clarify seeming contradictions in sex-specific physiology, such as the fact that females are more resilient to metabolic disease under some conditions while exhibiting increased disease risk in others^15,78,79^. Future work on molecular dimorphisms in adipocyte gene expression will provide new insights into how sex-specific adipocyte functions might underly disease vulnerabilities.

Sexually dimorphic physiology and behavior are instructed by genetic and transcriptional programs via downstream effectors of Tra, most of which are instructed by the transcription factors Dsx and Fru. In addition to expression differences between males and females, dimorphism in hormones such as 20E underly various aspects of sex-specific physiology. For example, recent work has demonstrated that female-biased plasticity of the adult intestine post-mating depends on high circulating 20E levels, which are higher in females than in males^80^. In larvae, female-biased expression of Ecdysone receptor (EcR), which regulates transcription of 20E-promoting gene targets upon binding to 20E, is essential for proper specification of the somatic gonad and maintenance of gametogenic potential in the adult^81^. EcR is expressed broadly in larval and adult tissues, including in the fat and ovary. EcR promotes a female metabolic state in the ovary by regulating expression of genes that support lipid biosynthesis and uptake^26^. ATF4 and EcR have been shown to physically interact *in vitro*^82^, raising the possibility of cooperation between ISR and EcR signaling pathways to promote a female metabolic state.

Metabolic pathways in lipogenic tissues like the fat and intestine drive many aspects of sex-specific physiology^1^. Research from over six decades ago establishes that in humans, a minimum requirement of fat stores within adipose tissue is required for onset of the fertile period in adolescence^83^. The prevailing thought is that evolutionary pressures led to a heavy reliance of the female reproductive system on metabolic tissues such as the fat and liver to produce metabolites/hormones^84^. In this paradigm, male metabolic tissues are less responsive to such evolutionary pressure due to the lower energetic cost of sperm production relative to egg production. The energetic “tradeoff” of this evolutionary pressure might underly consequent dimorphism in metabolic disease risk, wherein female metabolic tissues that have evolved to more efficiently store and utilize excess dietary fats show lower incidence of cardiometabolic disorders in comparison to males^7,14,78,79^. In extending our fundamental discoveries in this work to the clinical realm, an enticing postulation is that the dimorphism in ATF4 activity and/or adipose tissue physiology might contribute to the lower cardiometabolic risks reported in female humans. An important caveat that must be mentioned here is that not all stress resilience favors a female bias. For example, in a cohort of diabetic patients, females present a higher of developing cardiovascular disease^3^. Thus, future work in model organisms and humans will enable us to test the above postulation and its caveats.

### ATF4 induction in the fat enables adaptive response to nutrient deprivation

We found that higher ATF4 activity in females rendered them more resistant to metabolic stress imposed by *methioninase* expression (**Fig. 4; Fig. S4**). Curiously, despite higher basal levels of ATF4 activity in female adipocytes, we did not observe *Thor* induction to a higher degree in female fat body than in male fat body upon *methioninase* expression in comparison to adipocytes with control (*lacZ*) expression (**Fig. S4A**). This could reflect a limitation of our readout, which defines ATF4 activity quantitatively based on abundance of one known ATF4 target gene, *Thor* (*Drosophila 4E-BP*). While *Thor* induction is a widely employed readout for ATF4 activity, there may be other relevant target genes, yet unidentified, that are differentially expressed in males and females during nutrient deprivation stress. Future studies will examine transcriptional differences in male and female fat body during stress and degree of reliance on ATF4 activity.

Amino acid restriction has been shown to trigger ISR activation in *Drosophila* and ATF4 is known to be critical for mediating the response to such nutritional stress^48,51^. ISR activation is differentially sensitive to individual amino acids, though methionine deprivation robustly activates PERK- and GCN2-mediated ISR signaling^85^. Methioninase is bacteria-derived and catalyzes the conversion of sulfur-containing amino acids like methionine into α-keto acids, producing ammonia as a byproduct^86^. In the simplest interpretation, Methioninase expression results in methionine-deprivation, thus activating GCN2/ATF4 signaling. However, our study does not rule out the possibility that male-biased lethality upon *methioninase* over-expression is driven by the accumulation of α-ketomethionine, ammonia, or other downstream metabolites that result from processing of methionine. Interestingly, recent work has shown that elevated ammonia levels stimulate lipogenesis in the mammalian liver via ATF4 induction^87^, suggesting ammonia could be a minor contributor to the ATF4 induction we observe upon Methioninase induction.

### A potential role for ATF4 in regulating sexually dimorphic fat storage

Sexual dimorphism in fat storage mechanisms has been well documented^19,20,24–26^. Recent work by several groups^48,49,54,88,89^ has underscored the importance of ATF4 activity for homeostatic function, even in the absence of an obvious ‘stressor’ such as ER stress, dietary changes, immune challenge, or oxidative stress. One consensus from these recent studies underscores a role for ATF4 in lipid metabolism; loss of ATF4 is consistently found to lower overall stored fats in mice and *Drosophila*^49,54,55^. However, the total fat measurements in these studies did not include sex as a biological variable^90,91^, which is likely a major contributing factor to why the dimorphic nature of ATF4 signaling in fat tissues has previously gone unreported. We have recently shown that in the adult fat body, homeostatic ATF4 activity promotes the lipid droplet breakdown required for yolk synthesis^49^. Since lipid mobilization from larval fat body enables progression through metamorphosis^92^, our data support a model wherein sexually dimorphic ATF4 activity in larval adipocytes equips females with a higher capacity for lipid mobilization during this critical developmental period. Interestingly, sex differences in fat breakdown in adult *Drosophila* are influenced by dimorphic expression of transcripts encoding the TAG lipase, brummer (*bmm*), which is higher in adult male flies than in adult females^24^. We recently showed that the *bmm* locus contained several ATF4 binding sites, and a GFP reporter driven by these sites was de-repressed upon deletion of the binding sites^49^. Together, these observations lead us to hypothesize that dimorphic expression of lipid metabolism genes, potentially driven by ATF4, might underly sex differences in adipose tissue physiology. In addition, these principles could very well extend beyond the larval fat body, which we extensively test herein, to other dimorphic tissues that rely heavily on metabolic genes, such as the intestine and the brain.

In summary, our work establishes that part of the sexual differentiation program instructed by Tra/Dsx during development includes establishing higher basal ISR activity in female versus male adipocytes. Our findings indicate that increased resilience to nutritional stress in females can be ascribed, in part, to dimorphic ATF4 activity. Since ISR signaling is sensitive to changes in nutrient availability^56^, we propose that sexually dimorphic ISR signaling in *Drosophila* fat tissues may underly physiologically important molecular differences in metabolic dependency between males and females. Future work will investigate these differences at the molecular level, how they drive sex-biased stress resistance, and whether such processes govern metabolic sex differences and stress tolerance in mammals, including humans.

## Limitations of the study

### Specificity of GAL4 lines

Much of our results in this work rely on the use of *Dcg-GAL4* as a larval fat body driver, which enables robust transgene expression but is also expressed outside the fat body. Using *UAS-GFP* as a readout, we reliably observed *Dcg-GAL4* activity in both larval and adult adipocytes. However, we also found *Dcg-GAL4* activity in larval and adult hemocytes, which is consistent with previous reports^93^. The issue of tissue specificity is not limited to *Dcg-GAL4: R4-GAL4* was expressed in larval fat body, hemocytes, and gonads. In contrast, *3.1 Lsp2-GAL4* was faithfully fat-specific at all stages observed – absent from hemocytes, intestines, and gonads in both wandering L3 larvae and young adults – though its activity was low in larval tissues. *Lsp2* expression in the fat increases substantially in pupal stages such that *3.1 Lsp2-GAL4* activity is detectable in the adult fat body, consistent with previously reported *Lsp2* transcript abundance across developmental stages^94^. Despite these issues, we believe our primary results to be robust, though we cannot fully exclude the possibility of fat body non-autonomous roles in experiments that utilize *GAL4*-mediated transgene expression.

### Purity of larval fat body isolates for qPCR analyses

For all qPCR experiments in this manuscript, we dissected fat body from staged wandering third instar larvae and are careful to not include any other larval tissue. The exception to this is the larval gonad, which is embedded in the fat body. Based on *4EBP^intron^-GFP* reporter expression, we have found consistently that ATF4 activity is undetectable in larval gonads. Thus, we do not expect gonadal gene expression to be a significant contributor of the mRNA transcripts analyzed in this study. Since the gonad cannot be removed without considerably damaging the fat body, we elected to retain the gonad in our RNA preparations for qPCR. Indeed, our qPCR results demonstrating that ATF4 in the fat body is dimorphic is supported by visualization of enhancer-based reporters (**Fig. 1; Fig. S1**), they do not exclude the possibility that some of these sex differences in *crc* mRNA are due to differences in the gonad.

## Methods

### Fly stocks and husbandry

The transgenic and mutant lines used in this study are publicly available via Bloomington *Drosophila* Stock Center, Vienna *Drosophila* Resource Center, or were sourced from other labs. See **Table S1** for the complete list of lines used. *4EBP^intron^-GFP* transgenic flies were generated for this study; methods for construct design and transgenesis are described below.

Flies were reared on standard nutrient-rich agar medium containing cornmeal, molasses and yeast (LabExpress Inc., Ann Arbor, MI). All fly stocks were maintained at room temperature, and experimental crosses were reared in incubators at 24°C on a 12hr light/dark cycle.

### Immunostaining

Larval fat bodies were dissected in 1x PBS and fixed in 4% paraformaldehyde for 20 minutes at room temperature (RT) and washed twice in PBST (1xPBS + 0.1% Tween-20). For *Thor-lacZ* activity detection, samples were incubated overnight at 4°C with mouse anti-β-gal primary antibody (40-1a, 1:40, DSHB), followed by two washes in PBST and incubation with goat anti-mouse Alexa647 (1:500, Invitrogen) and DAPI (300 nM) for 20 mins in the dark at RT. For all other samples, fixation and the first two washes were immediately followed by DAPI incubation for 20 mins in the dark at RT. Samples were then washed twice in PBS and mounted in Vectashield (Fisher Scientific). Microscopy was performed using a Nikon A1 confocal microscope using a 20x objective lens.

### Generation of 4EBP^intron^-GFP transgenic animals

A 424-bp intronic element of the *Thor* (*4EBP*) locus, previously characterized to contain ATF4 binding sites (Kang 2017), was ligated into pStinger-attB via restriction cloning into SphI-XhoI sites. The resulting *4EBP^intron^-GFP* plasmid was then injected into *Drosophila melanogaster* embryos for stable genomic integration into the attP14 landing site (The Best Gene, Inc., Chino Hills, CA, USA). Primers used for cloning can be found in **Table S1**.

### Quantitative RT-PCR (qPCR)

Larval fat body was isolated from the posterior half of 3-4 animals per replicate for qPCR analyses. Number of animals per replicate/genotype was controlled within each experiment. Fat body preps included larval gonads and trivial amounts of Malphigian tubule tissue; thus, some of the RNA isolated for qPCR analysis represent contaminant transcript from these tissues. This is discussed further in the “Limitations of this study” section. α*Tub84B* was employed as a housekeeping gene for qPCR analyses, since levels do not vary significantly between males and females^95^.

### Developmental lethality assays

Assays were performed in triplicate at 24°C, with each assay vial containing 12 females and 3 males. To set up each assay, adults were anesthetized using CO_2_ and the correct number of females and males of each genotype were added to the assay vial. These assay parents were housed for 48 hours prior to assay to acclimate and recover from CO_2_ treatment. To perform the assay, parents were transferred by flipping into fresh vials containing nutrient-rich food (as described above). After 24 hours, parents were removed and flipped into new assay vials for a subsequent egg lay. For all egg lays, male parents were heterozygous for each listed allele, over a balancer. For example, control male genotype was *UAS-lacZ^RNAi^/CyO; UAS-lacZ/TM6B*. F1 adults of the correct genotype were then isolated and quantified for analyses. The cumulative male and female progeny that hatched from eggs laid during a 24-hour period were counted beginning 10 days following egg lay and for the following 5 days (to allow time for possible developmental delays). Animals were isolated at 0-2 days of age, at which time no lethality of eclosed adults was observed in any of the tested genotypes; this assay was not designed to capture adult lethality that might occur later than 2 days post-eclosion.

## Acknowledgements

We are grateful to the Center for Biological Imaging at the University of Pittsburgh for imaging assistance and access to equipment and to Dr. Andrey Parkhitko (University of Pittsburgh) for helpful discussion. Additionally, we thank the Bloomington *Drosophila* Stock Center (BDSC, Bloomington, IN), the Developmental Studies Hybridoma Bank (DSHB, Iowa City, IA), the Vienna *Drosophila* Resource Center (VDRC, Vienna, Austria), and the Transgenic RNAi Project (Harvard Medical School, Cambridge, MA) for making available reagents needed for this study. D.V. was funded by NIH R00EY029013 and NIH R35GM150516; L.G. was funded by NIH T32DK063922 and NIH K99GM149982.

## Author contributions

Conceptualization: L.G., D.V.; Methodology: L.G., D.V.; Investigation: L.G., M.M., D.V.; Formal analysis: L.G., M.M., A.A., D.V.; Validation: L.G., M.M., D.V.; Resources: L.G. and D.V.; Writing – original draft: L.G., D.V.; Writing – review and editing: L.G., M.M., A.A., and D.V.; Supervision: L.G., D.V.

---

**Figure S1.**
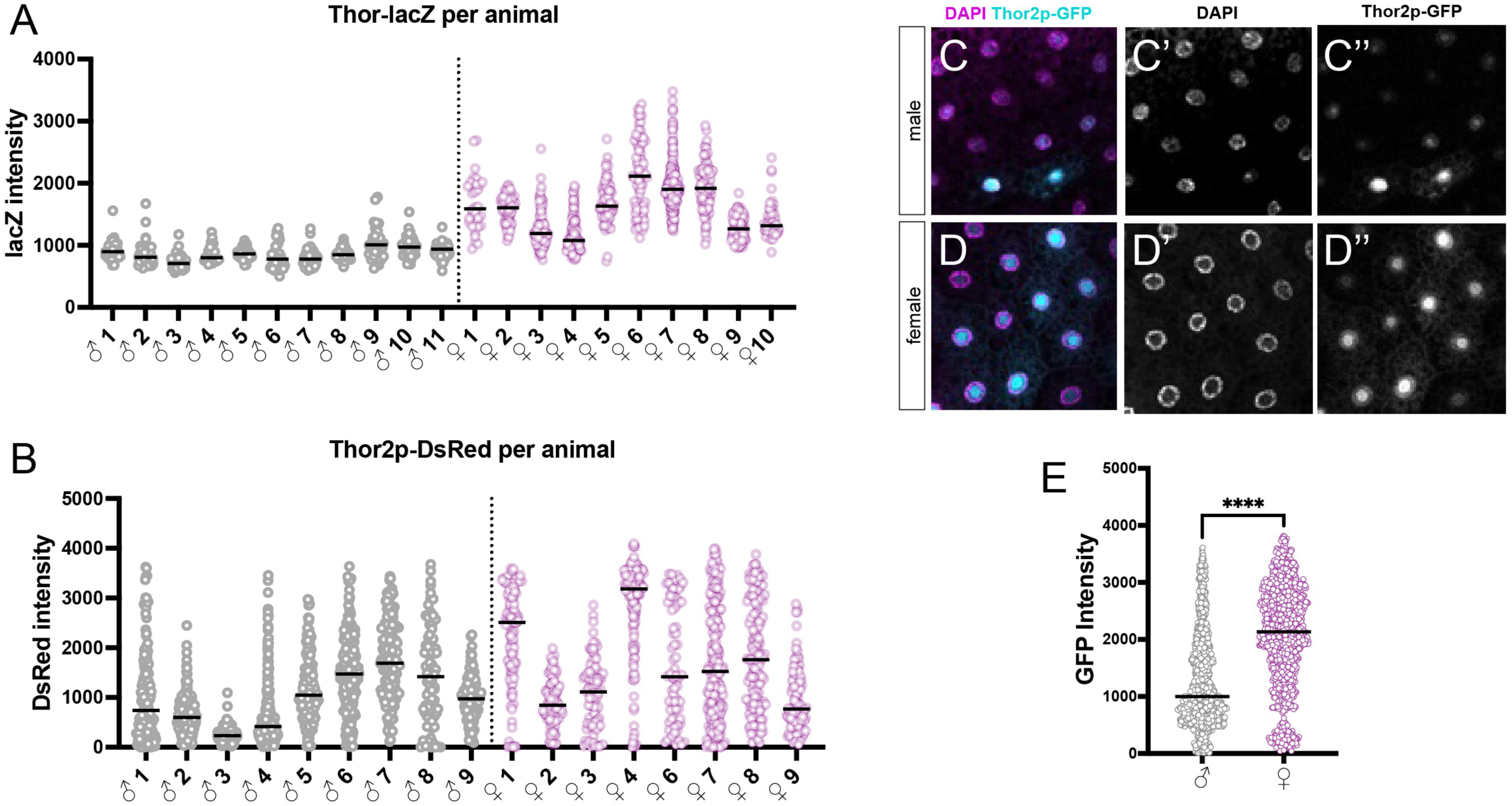
(A) Quantification of nuclear β-gal fluorescence intensity in adipocytes from individual *Thor-lacZ* male and female larvae to show animal-to-animal variability in reporter expression. (B) Quantification of nuclear DsRed fluorescence intensity in adipocytes from individual *4EBP^intron^-DsRed* male and female larvae to show animal-to-animal variability in reporter expression. (C-D) Representative immunofluorescence images of male (D) and female (E) larval adipocytes carrying a transgenic *4EBP^intron^-GFP* reporter. (E) Quantification of nuclear GFP fluorescence intensity in C-D.

**Figure S2.**
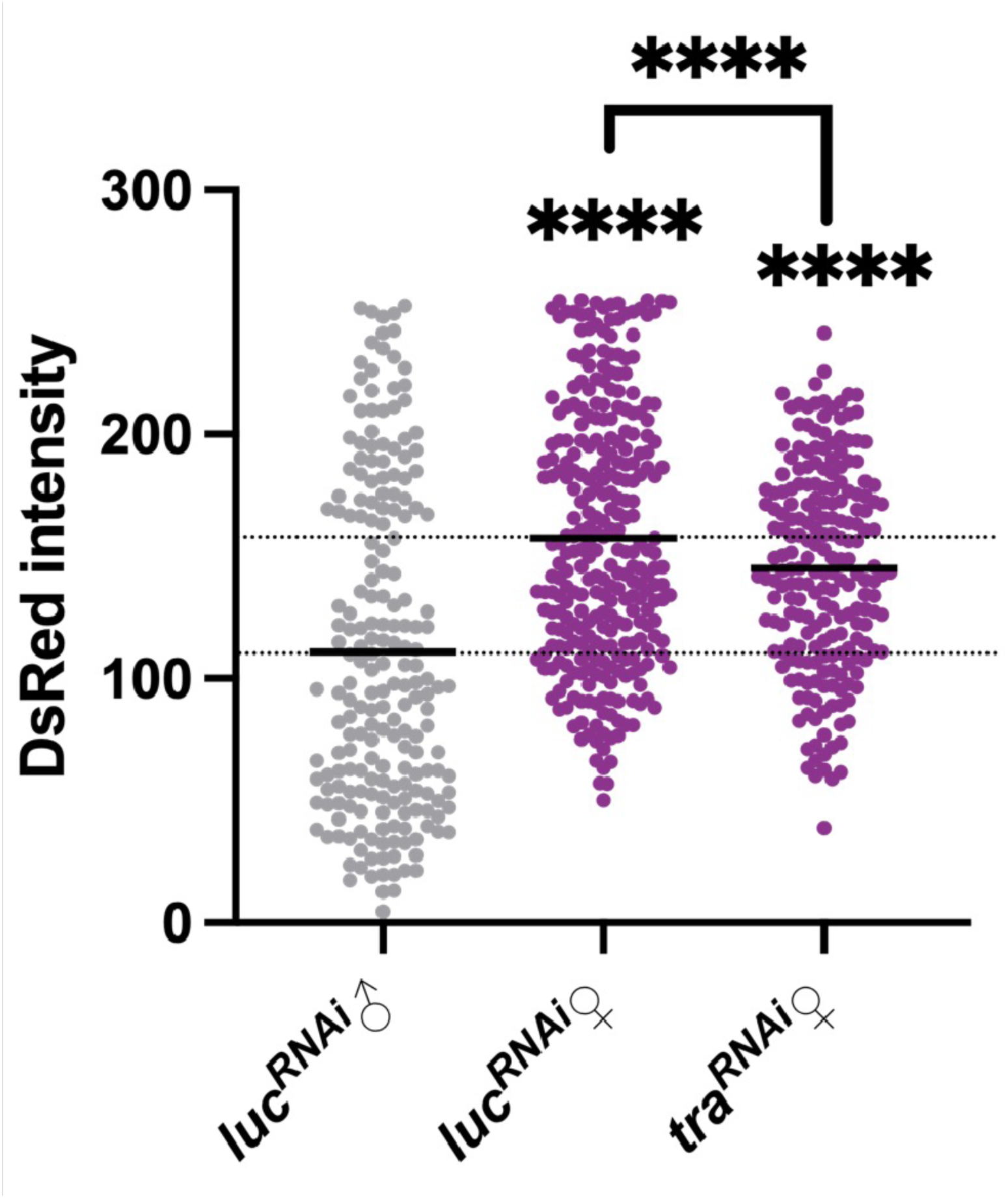
Quantification of nuclear fluorescence intensity of *4EBP^intron^-DsRed* reporter expression in control (*luc^RNAi^*) and genetically masculinized (*tra^RNAi^*) adipocytes.

**Figure S3.**
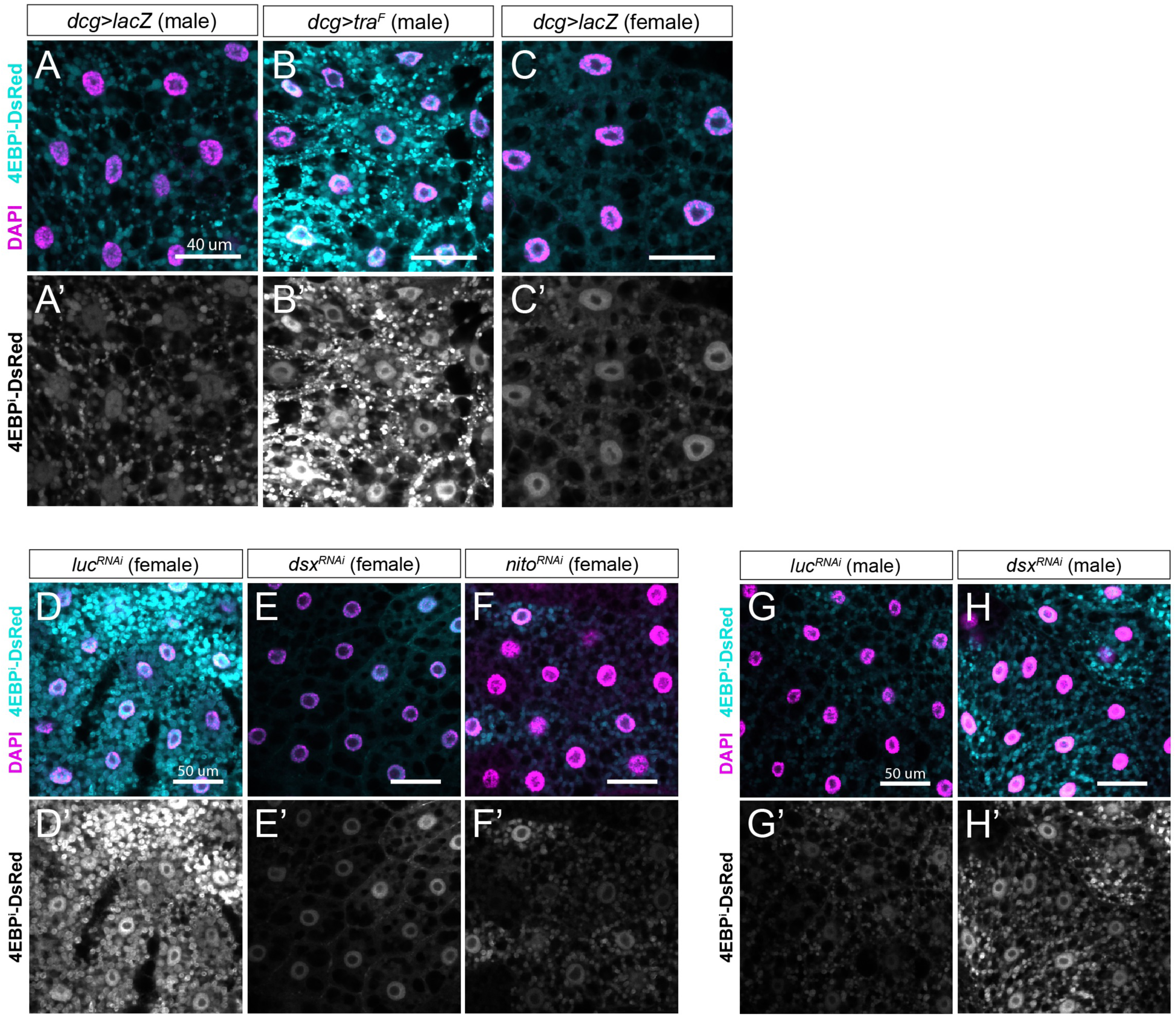
(A-C) Representative confocal images of control (*Dcg>lacZ,* A and C) or genetically feminized (*Dcg>tra^F^,* B) larval adipocytes. The demonstrated change in DsRed intensity is quantified in Fig. 3F. (D-F) Representative confocal images of *4EBP^intron^-DsRed* expression in adipocytes from female control (*Dcg>luc^RNAi^*, E) *Dcg>dsx^RNAi^* (F) or *Dcg>nito^RNAi^* (G) larvae. The observed changes in DsRed intensity are quantified in Fig. 3K. (G-H) Representative confocal images of *4EBP^intron^-DsRed* expression in adipocytes from male control (*Dcg>luc^RNAi^*, G) or *Dcg>dsx^RNAi^* (H) larvae. The observed changes in DsRed intensity are quantified in Fig. 3K.

**Figure S4.**
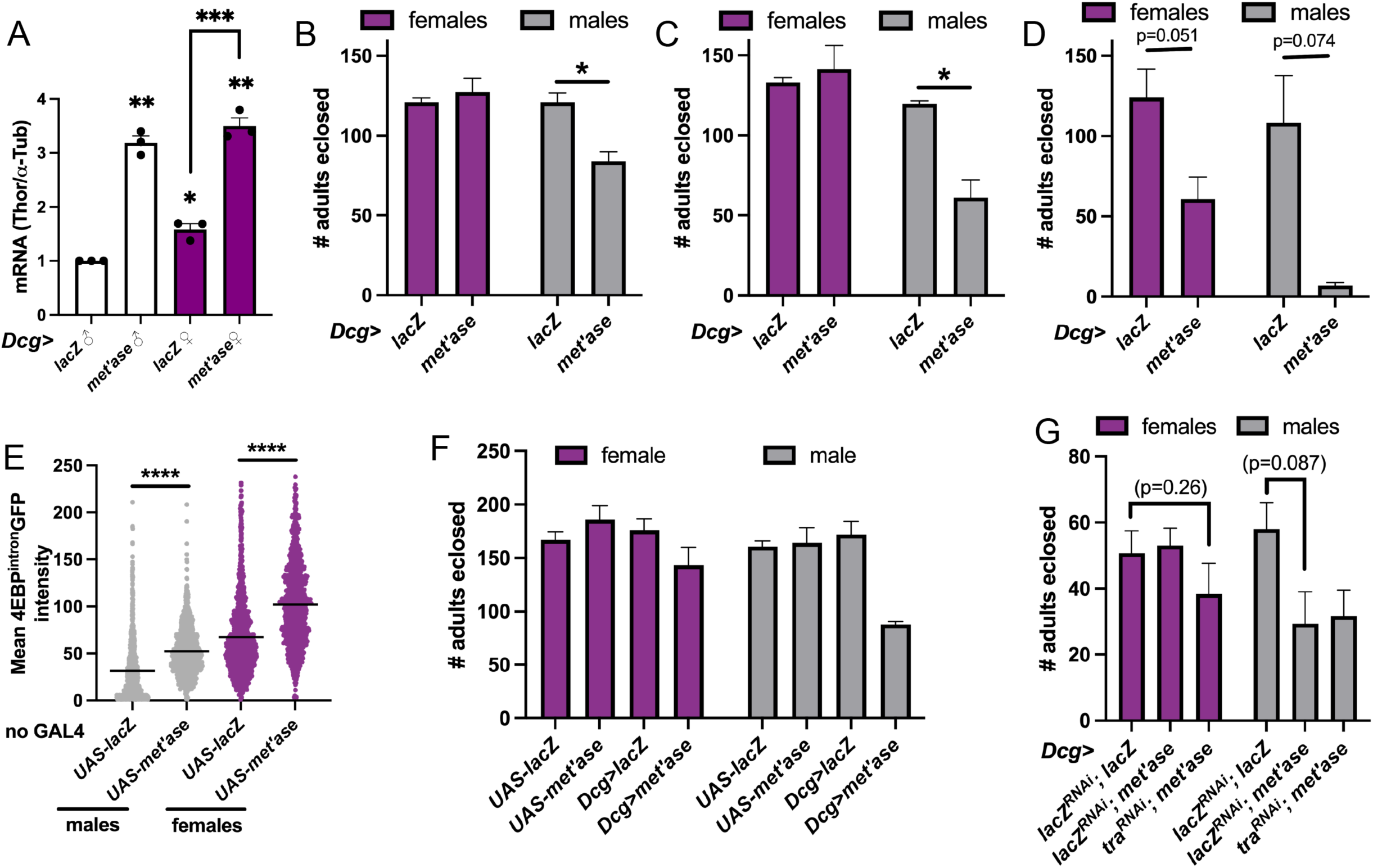
(A) qPCR analysis of Thor induction following *methioninase* expression in male and female larval adipocytes. *lacZ* expression is performed in control animals. (B-D) Quantification of adults eclosed following successive 24-hr egg lays, as performed in Fig. 4A, from *Dcg-GAL4* females crossed to either *UAS-lacZ (lacZ)* or *UAS-methioninase (met’ase)* males. Note that this graph is duplicated from Fig. 4A for ease of data interpretation. The same parents were used in **Fig. S4B-D** assays on consecutive days. Thus, female parental age was as follows: **S4B**: 5-7d; **S4C**: 6-8d; **S4D**: 7-9d. Male parental age was 0-2d in the first assay. (E) Quantification of nuclear *4EBP^intron^-GFP* fluorescence intensity in larval adipocytes heterozygous for either *UAS-lacZ* or *UAS-methioninase*, absent of a *GAL4* driver. Reporter expression was significantly higher in both male and female adipocytes carrying *UAS-methioninase* compared with control *UAS-lacZ* adipocytes, demonstrating that presence of the *UAS-methioninase* transgene alone is sufficient to trigger ATF4 induction. (F) Quantification of adults eclosed following successive 24-hr egg lays from *w^1118^* (first two bars of each color) or *Dcg-GAL4* females (last two bars of each color) crossed to either *UAS-lacZ* or *UAS-methioninase* males. Male lethality was observed in *Dcg>methioninase* animals (fourth gray bar), but not with *UAS-methioninase* crossed to *w^1118^* (second gray bar). (G) Quantification of adults eclosed following a 24-hr egg lay from *Dcg-GAL4* females crossed to males of the indicated genotypes (x-axis) to test protective role of female adipocyte sexual identity in preventing female lethality during *methioninase* expression. Statistical significance in A-D was evaluated using a series of unpaired Student’s t-tests with Welch’s correction.

**Table S1.** Transgenic and mutant flies used in this study.

| Genotype | Source | Figures used |
| --- | --- | --- |
| *P{lacW}Thor<sup>k13517</sup> (Thor-lacZ)* | BDSC #9558 | 1A-C, S1A |
| *4EBP<sup>intron</sup>-DsRed* | Dr. Hyung Don Ryoo<sup>51</sup> | 1D-F; 3A,F,K; S1B; S3 |
| *4EBP<sup>intron</sup>-GFP* | This study | S1C-E; 3A; S4E |
| *w<sup>1118</sup>* | BDSC #3605 | S2 |
| *Dcg-GAL4* | Dr. Jonathan Graff<sup>96</sup> | 3A-K; 4A,D-E; S3; S4A-D,F |
| *UAS-luciferase<sup>RNAi</sup> (TRiP JF01355)* | BDSC #31603 | 3A-E,K; S3D, G |
| *UAS-tra<sup>RNAi</sup>* (TRiP HMS02830) | BDSC #44109 | S2, 3B-E, S4G |
| *UAS-tra<sup>F</sup>* | BDSC #4590 | 3F-J, S3B |
| *UAS-lacZ* | BDSC #3956 | 3F-J,L; 4A-E; S3A, C; S4A-F |
| *UAS-methioninase* | Dr. Andrey Parkhitko<sup>67</sup> | 4A-E; S4A-F |
| *UAS-dsx<sup>RNAi</sup>* (TRiP HMC03795) | BDSC #55646 | 3K-L; S3E,H |
| *UAS-nito<sup>RNAi</sup>* (TRiP HMJ02081) | BDSC #56852 | 3K-L; S3F |
| *tra<sup>1</sup>* | BDSC #675 | 3A |
| *Tl{mCherry}tra<sup>KO.mCherry</sup> (tra<sup>KO</sup>)<sup>27</sup>* | BDSC #67412 | 3A |
| *R4-GAL4* | BDSC #33832 | 4B |
| *3.1 Lsp2-GAL4* | BDSC #84285 | 4C |
| *UAS-lacZ<sup>RNAi</sup>* | Dr. Hyung Don Ryoo | 4D |
| *UAS-Atf4<sup>RNAi</sup>* | VDRC #109014 | 4D |

**Table S2.**
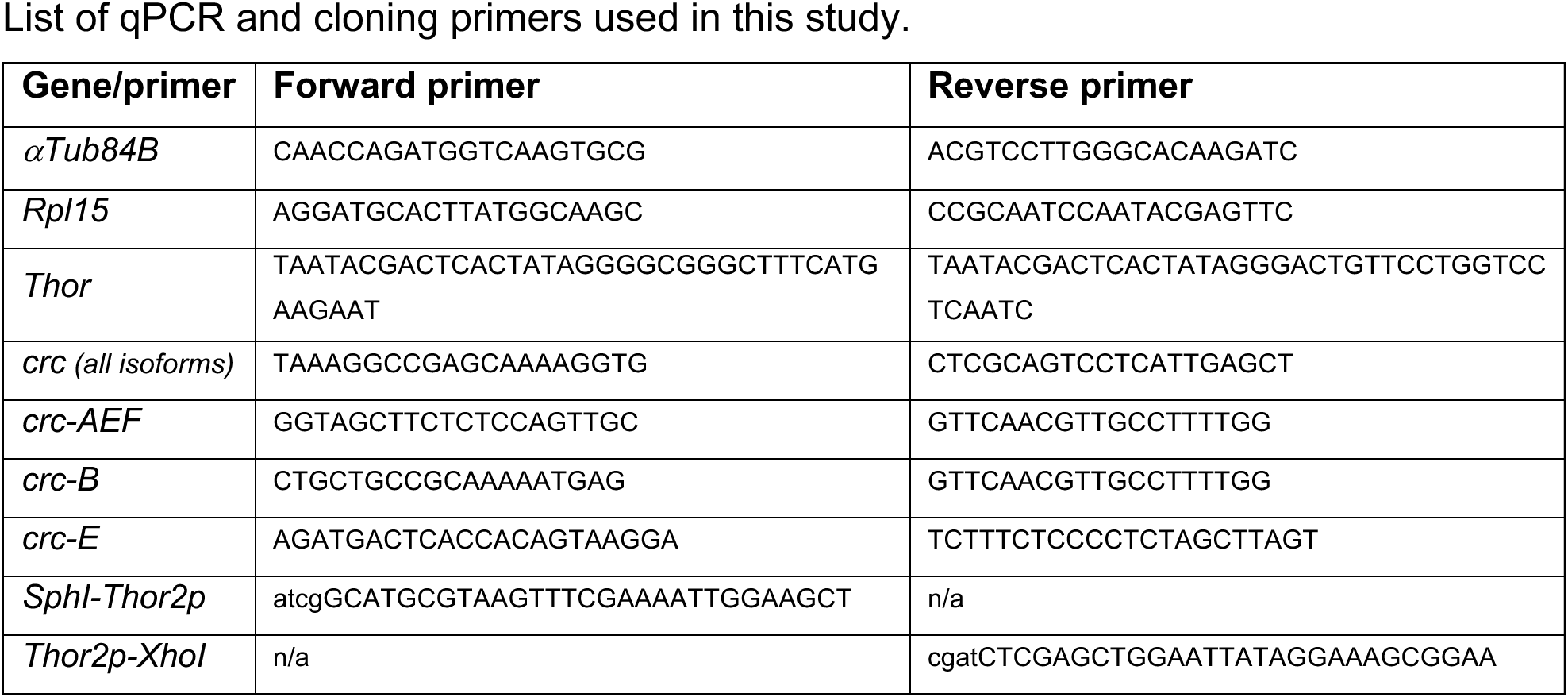
List of qPCR and cloning primers used in this study.

